# Spontaneous perceptual reversals are accompanied by systematic changes in pupil size but not respiration phase

**DOI:** 10.1101/2025.11.28.691130

**Authors:** Lisa Stetza, Christoph Kayser

## Abstract

Humans tend to align their respiration with important events, in line with the notion that respiration serves as a tool to allocate (neuro-)physiological resources. This respiration alignment is consistently reported in laboratory studies relying on sequences of stimulus-driven experimental trials. However, whether such respiration alignment also occurs in the absence of external cues - such as during spontaneous perceptual changes - remains unknown. However, pupil size is known to change around spontaneous changes in perception, and pupil size supposedly covaries with respiration phase, suggesting that indeed respiration may align to spontaneous changes in perception. We investigate the link between respiration, pupil size, and spontaneous perceptual changes while participants (n=21) reported spontaneous perceptual reversals while viewing an ambiguous Necker cube. We confirmed the biphasic modulation of pupil size around reversals (constriction followed by dilation) and the general variation of pupil size with respiration phase. However, this coupling of pupil size and respiration phase broke down around perceptual reversals. Crucially, our data reveal no clear alignment of respiration phase to perceptual reversals although respiration frequency was predictive of the perceptual stability duration. These findings suggest that the temporal predictability of external events is critical for the robust respiration alignment to cognitive events.

## Introduction

We tend to focus our respiration on critical events. In typical sensory-cognitive lab studies, for example, participants tend to align their respiration with the expected timing of individual trials and this alignment is predictive of how accurate or fast participants respond (Perl et al., 2019; Johannknecht & Kayser, 2022; Goheen et al., 2024; Harting et al., 2025; Stetza et al., 2025). At the same time, neural activity covaries with respiration. This includes a widespread modulation of brain activity with respiration at rest (Heck et al., 2017; Girin et al., 2021; Kluger & Gross, 2021) but also entails the covariation of neural activity directly relevant to the current task (Huijbers et al., 2014; Stetza et al., 2025; Kluger et al., 2021; Andrews et al., 2025; Tarrasó et al., 2025; Perl et al., 2019) and of activity reflecting processes related to arousal and attention with the respiration cycle (Andrews et al., 2025; Kluger et al., 2024; Kluger et al., 2021; Melnychuk et al., 2018; Melnychuk et al., 2021). This suggests that by aligning respiration with temporally expected sensory-motor events, the brain can ensure the optimal handling of these (Brændholt et al., 2023; Allen et al., 2022; Heck et al., 2017). Despite many studies supporting the alignment of respiration to external events it remains unclear whether an alignment of respiration to perceptual-motor events also extends to situations where external cues are absent, such as during spontaneous changes of (bistable) perception (Huijbers et al., 2014; Zelano et al., 2016; Arshamian et al., 2018; Nakamura et al., 2018; Perl et al., 2019; Park et al., 2020; Johannknecht & Kayser, 2022; Mizuhara & Nittono, 2023; Andrews et al. 2025; Harting et al., 2025; Saltafossi et al. 2025; Stetza et al., 2025).

Yet, such spontaneous changes in perception provide an important testbed to understand what factors drive the alignment of respiration. Previous studies on respiration alignment confound different putative drivers of this: one driver could be the expectation of specific externally timed requirements to act on sensory-motor contingencies, such as the expectation of an upcoming stimulus or the need to act on this (Kluger et al., 2021; Johannknecht & Kayser, 2022; Chalas et al., 2025). An alternative driver could be the intrinsic coupling of neural processes to respiration, by which the engagement of these processes in the experimental trial results in an alignment of respiration with the external event. Only in the latter case would one expect an alignment of respiration to spontaneous changes in perception, while the need for externally controlled expectations or sensory-motor contingencies would not predict an alignment to spontaneous changes in perception, as discussed in the following.

The apparent coupling of neural activity and respiration extends from sensory to association regions and most likely includes those regions involved in mediating spontaneous changes in perception, such as temporal, parietal and prefrontal cortex (Sterzer et al., 2009; Wang et al., 2013; Weilnhammer et al., 2017; Brascamp et al., 2018). Any modulation of neural processes in these regions along the respiration cycle should result in the spontaneous changes in perception to covary with the respiration phase, for example, by adjusting the gain of individual sensory representations or biasing decision processes. Further support for a potential alignment of respiration to spontaneous changes in perception comes from studies showing that pupil size changes around perceptual reversals (Einhäuser et al., 2008; Hupé et al., 2009; Brascamp et al., 2021; Nakano et al., 2021). Pupil size is a key marker of arousal (Aston-Jones & Cohen, 2005; Laeng et al., 2012; Viglione et al., 2023) and has been shown to systematically decrease prior to and increase after perceptual reversals (Brascamp et al., 2021; Nakano et al., 2021; de Hollander et al., 2018). At the same time, pupil size was shown to change systematically along the respiration cycle (Kluger et al., 2024; Schaefer et al., 2025), which may point to a common covariation of respiration with changes in arousal and spontaneous changes in perception.

Alternatively, respiration may align only to externally predicted events, be it the expectation of a particular sensory stimulus or the expectation to perform a specific motor behavior (Park et al., 2020; Park et al., 2022). In the absence of such events, the respiration alignment would vanish. The absence of alignment of respiration to spontaneous changes in perception could also be motivated by the observation that the time scale of neural activity underlying the encoding of typical external stimuli differs from that observed during intrinsic changes in perception. The neural responses evoked by external stimuli are typically characterised by a fast rise and a slow decay and usually emerge from primary sensory areas whose neural activity varies on a fast time scale (de Jong et al., 2020). In contrast, the neural activity associated with endogenous perceptual changes shows a more symmetric and slow rise and decay (Gelbard-Sagiv et al., 2018; de Jong et al., 2020; Wilson et al., 2023). Following this line of thought, one may not necessarily expect a relation between respiration and spontaneous changes in perception.

To address this question, we tested whether the respiration of human participants aligns to spontaneous changes in the perception of a bistable visual object. In our study, participants viewed an ambiguous Necker cube during prolonged periods (75-second trials) and reported their perceived perceptual reversals using a button press, while we recorded respiration and pupil size. We then tested whether respiration specifically aligns to perceptual reversals, whether reversals emerge during a particular respiration phase, and whether the duration of individual periods of perceptual stability is related to respiration phase or rate. In addition, we confirmed previous results showing that pupil size systematically changes around perceptual reversals and probed whether this covaries with the respiration phase, as suggested by previous work (Schaefer et al., 2025; Kluger et al., 2024).

## Methods

### Participants

A total of 32 adult volunteers participated in the study after providing written informed consent. All had normal or corrected-to-normal vision and hearing and were compensated for their time; participants consisted of young university students. Participants were informed about the experimental procedures and devices but were not explicitly told that the study focused on the relationship between respiration and task performance, similar to our previous work (Stetza et al., 2025; Harting et al., 2025; Johannknecht & Kayser, 2022). They were instructed to breathe normally through their nose, though we cannot exclude that some participants also breathed orally during the task. The Ethics Committee of Bielefeld University approved the study.

### Experimental setup and data acquisition

The experiment was conducted in an electrically shielded and sound-attenuated booth (Desone, Germany). Visual stimuli were presented on a computer monitor (27 inch; ASUS PG279Q, 120 Hz) positioned approximately 85 cm from the participant. Visual presentation was controlled using MATLAB (R2017a, MathWorks Inc., Natrick, MA) and the Psychtoolbox (Version 3.0.14). Synchronization of stimulus presentation and respiration recordings was implemented via TTL pulses sent to the BioSemi ActiveTwo system (ActiView software).

Respiration was recorded using a temperature-sensitive thermistor (Littelfuse GT102B1K, Mouser Electronics) attached to a modified single-use oxygen mask (Stetza et al., 2025; Harting et al., 2025; Johannknecht & Kayser, 2022). The changes in voltage generated by temperature fluctuations from nasal airflow during inhalation and exhalation were amplified and recorded via the analogue input of the BioSemi EEG system. Eye movements and pupil diameter were recorded monocularly using an EyeLink 1000 (SR Research Ltd., Canada) at 500 Hz in a free-viewing configuration. A 9-point calibration was performed prior to each block.

### Experimental task

Participants were first familiarized with the experimental task to ensure they understood and experienced spontaneous reversals of the Necker cube (11°x9°). For this, disambiguated variants of the cube with one face shaded in grey were presented to associate these with the respective response buttons (left and right arrow keys). During the actual task, participants completed 24 trials, each consisting of 75 seconds of continuous stimulus presentation, with their task being to report whenever the percept changed by pressing the corresponding button. Each trial was followed by a 15-second inter-trial interval displaying a uniform grey screen. The 24 trials were grouped equally into four blocks separated by breaks.

### Processing of respiratory data

Similar to our prior work (Stetza et al., 2025; Harting et al., 2025; Johannknecht & Kayser, 2022), the respiration signal was preprocessed using FieldTrip (Oostenveld et al., 2011) in MATLAB (R2022b). Signals were filtered using a third-order Butterworth filter (high-pass at 0.03 Hz, low-pass at 6 Hz), resampled at 100 Hz and converted to z-scores. The Hilbert transform was used to compute the analytic signal, from which local peaks were identified to define individual respiration cycles. Cycles were extracted in 7-second windows centred on these peaks, including only those peaks exceeding z = 0.5. Inspiration was defined based on a positive slope prior to a peak, and expiration based on a negative slope following this. In some cases, short pauses between inhalation and exhalation were classified as atypical data points and not assigned a specific phase (Noto et al. 2018). We also identified atypical respiration cycles by calculating the mean squared distance of all respiration cycles of each participant in time; cycles exceeding 3 standard deviations from the centroid were removed from data analysis. To link respiration to behavior the respiration phase was defined as a linear variable progressing from 0 to π during inspiration and π to 2π during expiration. This approach provides a continuous representation of the respiration phase with meaningful reference points (0/2π = peak inhalation; π = peak exhalation). We also determined the respiration frequency prior to each perceptual reversal based on the average of the instantaneous frequency of the last two cycles prior to each response.

### Processing of eye tracking data

The pupil diameter was extracted using custom MATLAB scripts and obtained following previous work (Brascamp et al., 2021): periods of missing data (blinks) were identified and linearly interpolated. The temporal drift of pupil data was corrected by fitting an exponential function to the data and subtracting this fit from the original signal. To reduce noise and artefacts, the detrended signal was low-pass filtered at 8 Hz. For alignment with respiration data, the filtered and smoothed signal was resampled to 100 Hz.

### Data exclusion and cleaning

The inspection of the behavioural data revealed a large inter-individual variability in the number of perceptual reversals, ranging from fewer than 120 to more than 1400 across trials. Such variability can complicate data analysis and the comparison with previous studies. To standardize the dataset we excluded participants whose reversal count fell outside an empirically defined range: given a total task duration of 1800 seconds, we applied an upper cutoff of 900 reversal (i.e., one every 2 seconds), accounting for the fact that some neuroimaging studies using bistable stimuli or binocular rivalry have reported neural precursors 1-2 seconds prior to reversals (Wilson et al., 2023; Gelbard-Sagiv et al., 2018). We also required a minimum of 120 reversals to obtain sufficient data epochs for analysis. Based on these criteria, we excluded data from 10 participants from further analysis (6 exceeded the maximum, 4 fell below the minimum). For some participants the eye tracking did not work reliably during some of the experimental blocks. For the final analysis we hence stratified the dataset to include only those participants with robust behavioral data and reliable eye tracking in most of the trials. This resulted in a sample of N = 21 participants. For the analyses linking respiration and behavior, we included on average 333.2 ± 161.2 (mean ± SD) trials per participant. For the analyses linking respiration and pupil size, we included only trials with reliable eye tracking data, which resulted in 285.9 ± 145.6 (mean ± SD) trials per participant.

### Alignment of respiration and pupil size to perceptual reversals

Previous studies have demonstrated the alignment between respiration phase and task events using a metric of phase coherence (Goheen et al., 2024; Johannknecht & Kayser, 2022; Kluger et al., 2021; Stetza et al., 2025; Harting et al., 2025; Saltafossi et al., 2025). We here used this approach to quantify the respiration alignment to perceptual reversals within participants based on the respiration data (sampled at 100 Hz) in time epochs of 4 seconds around each indicated reversal. We converted the respiration phase to a complex number and computed the vector length of the average of interest (Park et al., 2020; Johannknecht & Kayser, 2022): to obtain the within-participant coherence, we averaged across all epochs within a participant. The statistical significance of the resulting coherence values (the group-average of the epoch-by-epoch coherence) was tested against a null distribution generated from 4000 random temporal shifts of respiration phase under the assumption of no alignment to the experimental task (Stetza et al., 2025; Harting et al., 2025; Johannknecht & Kayser, 2022; Kluger & Gross, 2021). For each randomization, we derived the maximal value over time points to correct for multiple comparisons over time. Additionally, we tested for a correlation between the within-participant phase coherence (at the time of maximum coherence in the across-participant averaged phase coherence) and the number of reported perceptual switches, using the original reversal number before data cleaning. For pupil size, we first computed the epoch-averaged pupil trace for each participant and then derived the group-average of these. We then used a cluster-based permutation test to test the null hypothesis of whether the group-average differs from zero (i.e., baseline)(Nichols & Holmes, 2002; Maris & Oostenveld, 2007). The first-level inference was derived based on point-wise t-tests of the pupil size to differ from zero and these first-level effects were thresholded at p<0.05. Significant time bins were clustered based on a minimal cluster size of two and using the maximum as cluster statistics. We then contrasted clusters in the actual data with clusters in a surrogate distribution obtained from 20000 permutations of the first level effects. We report significant clusters at a second-level significance of p < 0.01.

### Analysis of the predictive power of respiration and pupil size on percept stability

We used linear mixed-effect models to probe whether the phase of respiration, the rate of respiration, or pupil size holds predictive power for the stability of the individual percepts. We tested this both for the stability of the percept prior to each indicated reversal (i.e., the duration between the current and previous reversal) and the stability of the following percept (i.e., the duration between the current and next reversal). Stability durations were normalized within participants by their respective average to account for the between participant variability. The phase of respiration was coded as the sine and cosine components of the cyclical phase value. We used a framework of model comparison to establish the predictive power of each factor of interest, similar to our previous studies (Johannknecht & Kayser, 2022; Harting et al., 2025; Stetza et al., 2025). For the respiration phase, we contrasted two models, one including respiration and one without:

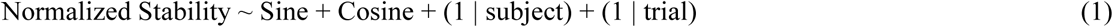

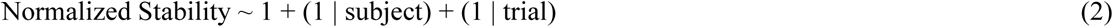

We then derived the Akaike Information criterion (AIC) for each model and used their difference for model comparison. From these, we derived the conditional probability of each model given the data, which can be obtained using associated Akaike weights (Wagenmakers & Farrell, 2004). We considered an AIC difference > 9.2 as evidence for an effect of respiration, which corresponds to a probability of above 99% that the model including respiration has more explanatory power than the alternative model of above 99%. We performed this model comparison for the respiration phase at multiple time points in the epoch around each reversal, sampled at 200ms.

Using the same approach, we quantified the predictive power of 1) the respiration frequency on the stability of the individual percepts, 2) pupil size on the stability of the individual percepts, and 3) respiration phase on pupil size.

## Results

For the 21 participants included in the main analyses, the average reversal rate was 4.38 ± 3.49 (mean ± SD) seconds. The average duration of respiration cycles was 3.78 ± 0.72 seconds (mean ± SD), corresponding to a respiration rate of (0.27 ± 0.05 Hz). Inspiration phases were slightly shorter (1.71 ± 0.43 s) than expiration phases (2.09 ± 0.43 s).

### Respiration is only weakly aligned to perceptual reversals

To test whether participants’ respiration phase was systematically aligned to perceptual reversals, we computed the epoch-by-epoch coherence of the respiration phase for each participant. This within-participant phase coherence did not exceed chance level at p<0.01 (Fig. 2A; black dotted line, corrected for multiple comparisons along time) but did reach significance from -1.95 to -1.44 and from 0.71 to 1.51 seconds at p<0.05 (Figure 2A, grey dotted line). The observed coherence values (peak = 0.117) were also much smaller compared to those obtained in previous studies (phase coherence values between 0.2-0.25) testing the alignment to external sensory events (Harting et al., 2025; Kayser et al., 2025; Stetza et al., 2025). Figure 2B shows the individual phase coherence values at the time of group-average maximum (1.1 seconds). The participant-wise epoch-averaged phase (Figure 2C) also reveals considerable heterogeneity across participants.

**Figure 1:**
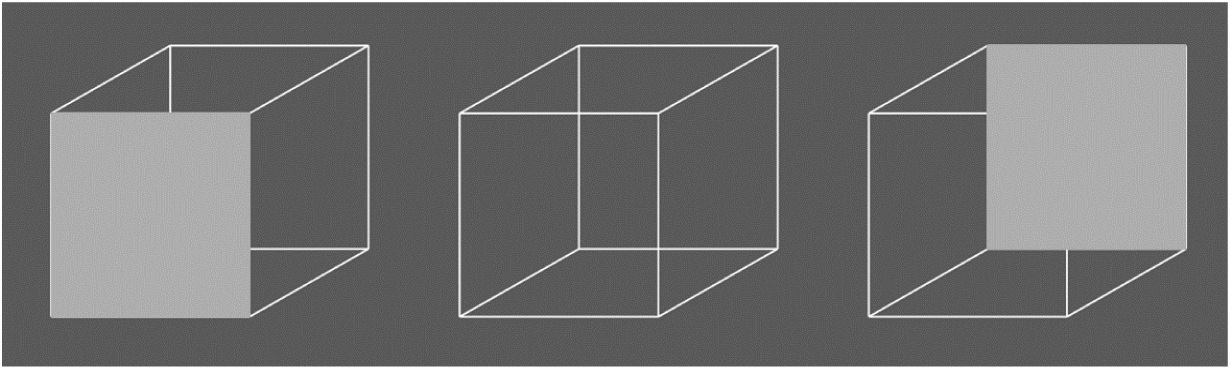
Visual stimulus. The Necker cube was displayed to participants to associate the two response keys with each of the two interpretations of the cube (left and right cubes). The middle panel illustrates the ambiguous cube presented during the actual experiment.

**Figure 2:**
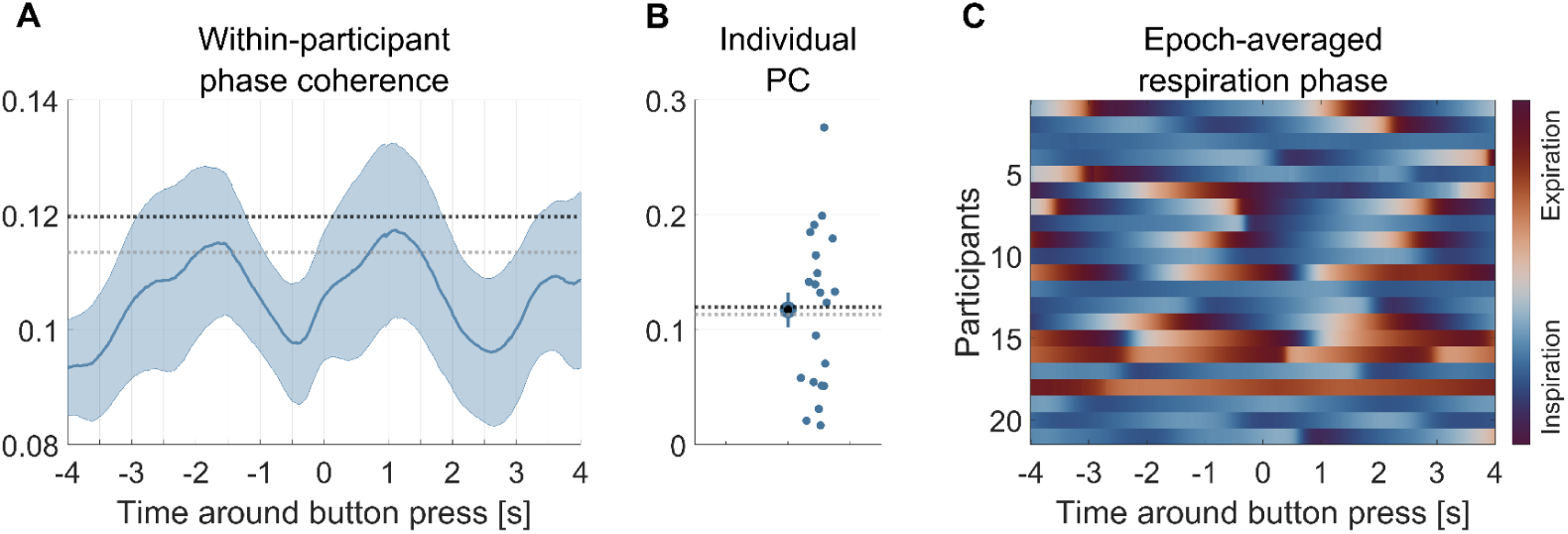
Respiration phase around reversals. A) Within participant coherence of the respiration phase (mean ± SEM). Dotted lines indicate significance level at p<0.01 (black) and p<0.05 (grey). Coherence did exceed p<0.05 level from -1.95 to -1.44 and 0.71 to 1.51 seconds. B) Participant-level phase coherence values at the time point of maximum phase coherence in the group average displayed in panel A (max phase coherence = 0.11 at 1.1 seconds relative to reversal). Dotted lines as in panel A. C) Epoch-averaged respiration phase for each participant. N = 21.

The correlation of the total reversal number and within-participant phase coherence showed a negative relationship, with individuals reporting lesser reversal showing higher coherence in their respiration alignment, but this correlation did not reach significance (r = -0.358, p = 0.111).

### Respiration frequency but not respiration phase explains percept stability

We then asked whether the respiration phase at the time of button press is predictive of the duration of the percept preceding or following the respective reversal. To test this, we compared the predictive power of linear models including or omitting the predictor of interest. Across the tested time points, the AIC differences did not reach significance for either perceptual duration (Figure 3A; preceding percept: max deltaAIC = 2.87 at 4 s, p = 0.808; following percept: max deltaAIC = 5.068 at -2.2 s, p = 0.812). Figure 3B displays the individual for the following percept, showing the (within-participant normalized) duration of the percept against the binned respiration phase at the time of the indicated reversal.

**Figure 3:**
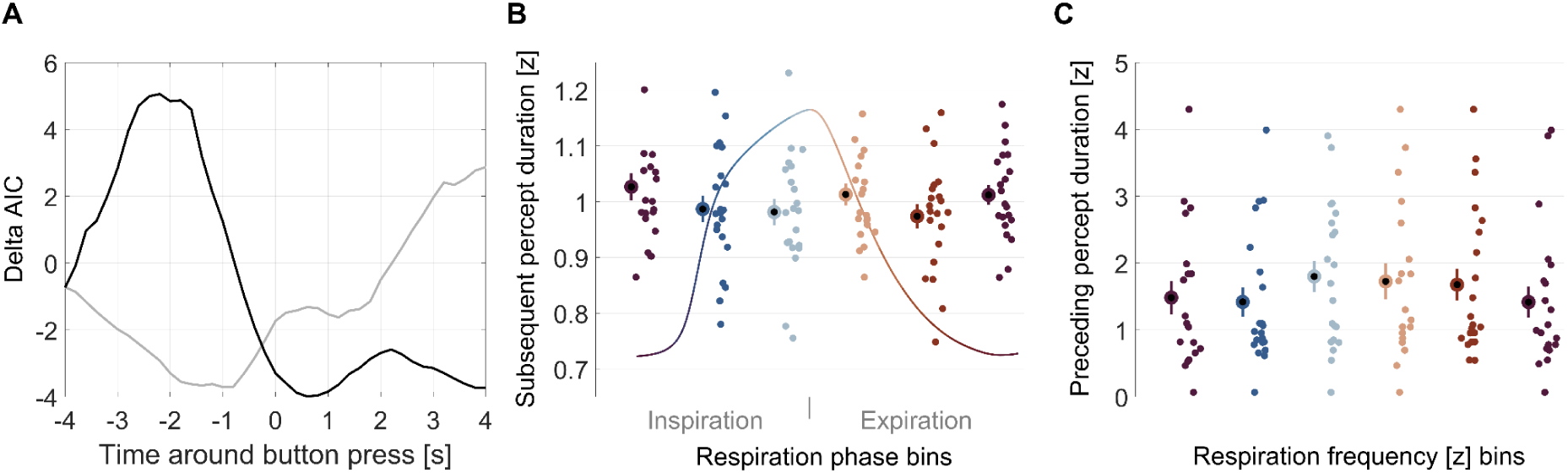
Respiration phase and frequency do not predict percept stability. A) Delta AIC values for models predicting normalized percept duration based on the respiration phase at different time points in an epoch ± 4 seconds around perceptual reversals compared to a null model. Grey: predicting the duration of the preceding percept, max deltaAIC = 2.87 at + 4 seconds; black: predicting the duration of the subsequent percept, max deltaAIC = 5.06 at -2.2 seconds. B) Normalized duration of the subsequent percept as a function of binned respiration phase. C) Normalized duration of the preceding percept as a function of the (normalized) respiration frequency. Bold dots show the group average with errorbars showing the SEM, smaller dots represent individual participants.

We then tested whether the respiration frequency is predictive of the duration of the percepts. We found that this is indeed the case: respiration frequency had significant predictive power on both the durations prior and following the reversals (deltaAIC pre-duration = 323.9, p<10^-5^; deltaAIC post-duration = 393.1, p<10^-5^). Investigating the respective model parameters revealed that the model for the preceding reversal yielded a significant negative relationship, indicating that with higher respiration frequency the percept stability shortens (slope = -0.022, t = -2.178, p = 0.029). Figure 3C illustrates the respective effect. For the subsequent precept the slope of the respective model was not statistically significant (slope = 0.001, t = 0.111, p = 0.912).

### Pupil size changes around reversals but is not predictive of perceptual stability

Pupil size systematically changed around the perceptual reversals as reported in previous work (Brascamp et al., 2021; Nakano et al., 2021; de Hollander et al., 2018): the pupil constricts preceding the reversal and dilates shortly after the indicated reversal, as shown in Figure 4A (cluster-based permutation test, one cluster from -1.58 to 0.05 seconds, p= 0.0141; one cluster from 0.34 to 1.57 seconds, p= 0.0106). Similar as for respiration, we then asked whether pupil size is predictive of the stability of the perceptual states. However, this was not the case, neither for the perceptual duration prior to the reversal (max deltaAIC = 7.69 at 1 s, p = 0.979) nor for that following the reversal (max deltaAIC = 6.10 at 0.6 s, p = 0.955).

**Figure 4:**
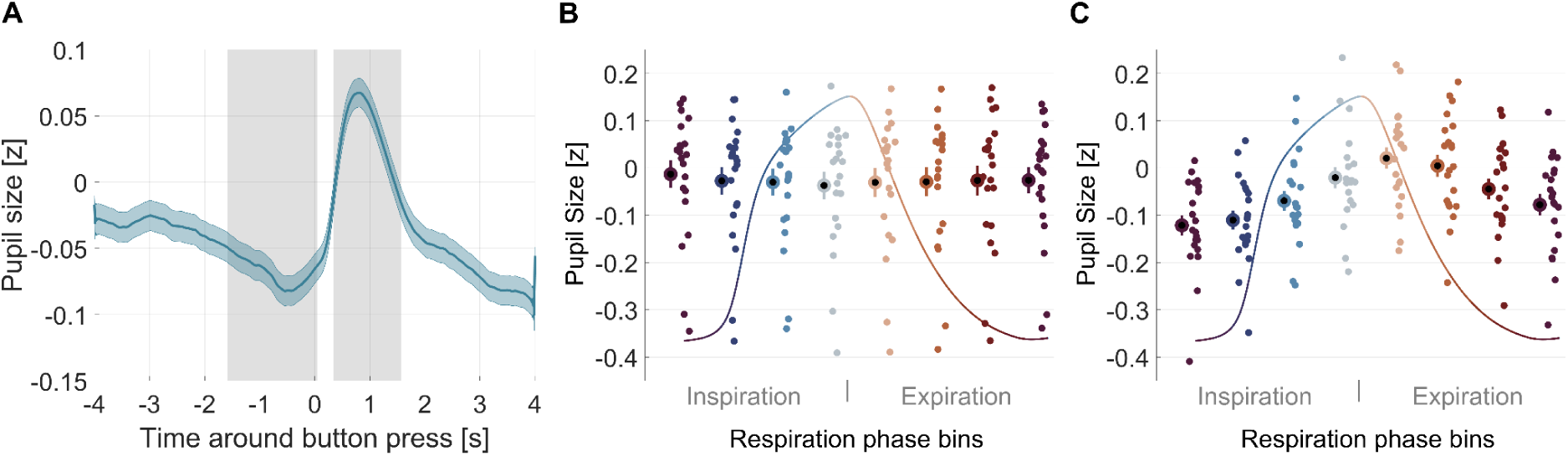
**Pupil size around reversals**. A) Group-averaged pupil trace around perceptual reversals (mean ± SEM). Grey shaded areas show epochs in which the pupil size differs significantly from zero (cluster-based permutation test, one cluster from -1.58 to 0.05 seconds, p= 0.0141; one cluster from 0.34 to 1.57 seconds, p= 0.0106). B) Pupil size as a function of respiration phase (8 bins) in the epoch around reversal. C) Pupil size as a function of respiration phase for all data points in the entire 75-second trials. Bold dots show the group average with errorbars showing the SEM, smaller dots represent individual participants.

### Pupil size changes with respiration phase overall but not during reversal epochs

Finally, we asked if the respiration phase is predictive of pupil size, as suggested by previous work. We first tested this within the epochs around the reported reversals. This revealed no significant predictive power of respiration phase (max delta AIC = 0.893 at -3.8 s, p = 0.610, Figure 4B). We then tested this based on the data at all time points during the entire experimental trials, hence including periods of reversals and periods of longer perceptual stability. This revealed a significant predictive power of respiration phase (delta AIC value = 470.65, p<10^-5^), as reported in previous work (Schaefer et al. 2025; Kluger et al. 2024). Figure 4C shows the relationship between pupil size along the respiration cycle, with the pupil being smallest around inspiration onset and largest around expiration onset.

## Discussion

The present study examined the three-fold relation between respiration, pupil size and spontaneous changes in perception. Our data reproduce the previously reported alignment of pupil size to perceptual reversals and the general covariation of pupil size with respiration phase. However, our data suggest only a weak, if existent, alignment of respiration phase to perceptual reversal. The respiration phase was also not predictive of the stability of the perceptual periods, although respiration frequency was. Overall, this suggests that there is no systematic and strong relation of respiration phase with spontaneous changes in perception, unlike in externally driven sensory-cognitive tasks where respiration predictively aligns with upcoming experimental trials.

### Respiration phase is only weakly aligned to perceptual reversals

While previous studies using externally-driven sensory-cognitive tasks report a robust alignment of respiration phase to individual trials (Flexman et al., 1974; Gallego et al., 1991; Huijbers et al., 2014; Zelano et al., 2016; Arshamian et al., 2018; Nakamura et al., 2018; Perl et al., 2019; Park et al., 2020; Johannknecht & Kayser, 2022; Mizuhara & Nittono, 2023; Andrews et al. 2025; Braendholt et al., 2025; Harting et al., 2025; Saltafossi et al. 2025; Stetza et al., 2025), endogenously produced changes in perception seem to lack such a clear relation. For the present paradigm, the peak phase coherence was weak when compared to the phase coherence reported in previous studies that used external stimuli (Harting et al., 2025; Kayser et al., 2025; Stetza et al., 2025; Goheen et al., 2024), and reached only a moderate level of significance. Importantly, in sensory-cognitive tasks, the alignment seems to emerge 2-3 seconds prior to individual trials, suggesting that respiration actively aligns to the expected events. In the present paradigm an external, predictable occurrence of external stimuli is absent. Hence, the weak respiration alignment observed here suggests that the anticipation of an upcoming stimulus may be a potential key driver for the respiration alignment. Indeed, emerging work suggests that precisely this external temporally predictive information is critical to facilitate respiration alignment, as both spatial cues and temporal cues were shown to facilitate respiration alignment and increase perceptual sensitivity in a visual detection task (Chalas et al., 2025). Our results corroborate this notion of temporal prediction as a key driver of alignment, suggesting that external guiding factors facilitate and thereby enable more coherent respiration alignment.

### Respiration frequency might coordinate perceptual reversals in the absence of external references

While respiration does not seem to be specifically aligned to the indicated perceptual reversals, it is possible that perception in general is still entrained by the rhythm of respiration. In support of this, we observed the strong predictive power of respiration frequency on the duration of the perceptual stability. This suggests that the overall pace of respiration and that of perceptual switches are still related. While the present data cannot disentangle the causal relationship, whether respiration frequency paces switches in perception, or vice versa, this aspect of our data may point to a profound link between cycles in perception and cycles in respiration. However, the variability of the perceptual durations in typical data on bistable perception tends to follow a highly skewed and wide distribution (Wilson et al., 2023; Wang et al., 2013; Nakano et al., 2021), while the variability in the cycle durations of respiration is much smaller. While this does not invalidate a putative coupling of perceptual and respiration cycles, it also suggests that any such relation is not straightforward. Future studies employing manipulations of participants’ respiration patterns, or of manipulations of the stability of bistable percepts (Maier et al., 2003; Leopold et al., 2002; Brascamp et al., 2009), could be used to investigate this relation between respiration frequency and perceptual cycles further.

### Temporal dynamics of perceptual reversals may limit respiration alignment

It could also be that the alignment of respiration to perceptual reversals is shadowed by the temporal variability and sluggishness of the underlying neural processes. Neuroimaging studies support that paradigms involving bistable perception engage higher-order brain networks comprising frontoparietal and temporal areas (Sterzer et al., 2009; Wang et al., 2013). These networks reflect the integration of feedforward and feedback signals (Weilnhammer et al., 2017) and meta-analytic studies support the view that reversals arise from gradual, distributed interactions across multiple hierarchical levels (Brascamp et al., 2018). Neural activity in those regions evolves over slower timescales than the activity in early sensory regions (Gelbard-Sagiv et al., 2018; Wilson et al., 2023; de Jong et al., 2020). Hence, perceptual reversals are accompanied by widespread and relatively slow and gradual transitions of brain activity, rather than brief and immediate changes that are often associated with sudden external signals (de Jong et al., 2020). For example, some studies reported differences in EEG activity between ambiguous and unambiguous stimuli emerging around 1 second prior to reversals, and activity related to the percept of binocular rivalry stimuli as early as 2 seconds prior to the reports (Wilson et al., 2023; Gelbard-Sagiv et al., 2018). Perhaps then the slow buildup and variability of neural transitions underlying perceptual reversals lack the sharp and reliable onset necessary for reliable respiration alignment, despite the widespread covariation of respiration and brain activity across sensory and association cortices (Kluger & Gross, 2021; Goheen et al., 2023).

### Pupil size changes systematically around reversals and is linked to respiration phase

We observed a consistent modulation of pupil size locked to perceptual reversals, with a constriction preceding the reported changes in perception and a pronounced dilation following this. This bi-phasic change in pupil size is in line with previous studies. Following previous studies, the pupil constriction prior to a reversal may reflect the action of the noradrenergic arousal system on neural representations of the different interpretations of the bistable image (Hupé et al., 2009; Brascamp et al., 2021; Nakano et al., 2021). The pupil dilation following a reversal, in contrast, may in part also reflect task requirements and motor-related processes (Einhäuser et al., 2008; Hupé et al., 2009; Brascamp et al., 2021; Nakano et al., 2021).

In our data the trial-wise pupil size was not predictive of the persistence of the individual percepts prior to a reversal or those following this. That is, while our data confirm a general biphasic change in pupil size around reversals, previous studies also reported a relationship between pupil size and the persistence of individual percepts (de Hollander et al., 2018; Brascamp et al., 2021; Einhäuser et al., 2008). For example, one study reported that the pupil size prior to a reversal is predictive of the persistence of the following percept (de Hollander et al. 2018), while another study reported the covariation of pupil size prior to a reversal with the persistence of the current percept (Brascamp et al. 2021). Hence, previous work also remains somewhat inconclusive about the directionality of a putative relationship between pupil size and the temporal persistence of sensory percepts. One potential explanation for why we did not find such a dependency may also lie in the durations of the individual percepts, which in the present data were considerably shorter than in some previous studies (Wang et al., 2013; Nakano et al., 2021; Wilson et al., 2023). Future studies are required to corroborate a link between pupil size, arousal and the stability of individual percepts.

In line with previous studies, we found that pupil size was significantly related to respiration phase when considering the entire data across the experimental trials. Such a covariation of pupil size and respiration phase has been reported during resting periods and cognitive demanding periods in previous studies (Kluger et al., 2024; Schaefer et al., 2025; Melnychuk et al., 2018). This may reflect the differential engagement of arousal along the respiration phase, presumably enabled by the bidirectional projections between the locus coeruleus and the preBötzinger complex (Yackle et al., 2017). One possibility is that variations in arousal arising from variations in task performance are relayed to the preBötzinger complex and thereby directly shape respiration according to the current task demands. However, when restricting the analysis to the 4-second epochs around perceptual reversal, we did not find a significant relation between pupil size and respiration phase. This suggests that the overall covariation of both signals breaks down during moments related to specific epochs of the task, similar to what one previous study reports using a delayed matching-to-sample task (Nakamura et al., 2019). It could, for example, be that a more rapid change in arousal preceding a change in perception is too fast to enable a concomitant change in respiration phase, leading to a dissociation in the two signals. Alternatively, or in addition, neural processes related to the motor action involved in indicating the perceived reversal could influence arousal and pupil size, again acting on a time scale too fast to affect respiration. A final possibility is that the coupling of pupil and respiration seen here in general, and in other sensory-cognitive tasks in previous work, relates to a common entrainment of both signals by the temporally regular and predictive structure of the task context. Since this is lacking here, at least on the precise time scale of individual reversals, this may contribute to the dissociation of the two signals. Clearly, future work is required to address these points in more detail.

## Author contributions

L.S. and C.K. designed research; L.S. performed research; C.K. contributed unpublished reagents/analytic tools; L.S. and C.K. analyzed data; L.S. and C.K. wrote the paper.

## Additional information

The authors declare no competing financial and non-financial interests.

